# Ring formation by *Vibrio* fusion protein composed of FliF and FliG, MS-ring and C-ring component of bacterial flagellar motor in membrane

**DOI:** 10.1101/2023.02.06.527414

**Authors:** Kanji Takahashi, Tatsuro Nishikino, Hiroki Kajino, Seiji Kojima, Takayuki Uchihashi, Michio Homma

## Abstract

The marine bacterium *Vibrio alginolyticus* has a single flagellum as a locomotory organ at the cell pole, which is rotated by the Na^+^-motive force to swim in a liquid. The base of the flagella has a motor composed of a stator and rotor, which serves as a power engine to generate torque through the rotor–stator interaction coupled to Na^+^ influx through the stator channel. The MS-ring, which is embedded in the membrane at the base of the flagella as part of the rotor, is the initial structure required for flagellum assembly. It comprises 34 molecules of the two-transmembrane protein FliF. FliG, FliM, and FliN form a C-ring just below the MS-ring. FliG is an important rotor protein that interacts with the stator PomA and directly contributes to force generation. We previously found that FliG promotes MS-ring formation in *E. coli*. In the present study, we constructed a *fliF*–*fliG* fusion gene, which encodes an approximately 100 kDa protein, and the successfully production of this protein effectively formed the MS-ring in *E. coli* cells. We observed fuzzy structures around the ring using either electron microscopy or high-speed atomic force microscopy (HS-AFM), suggesting that FliM and FliN are necessary for the formation of a stable ring structure. The HS-AFM movies revealed flexible movements at the FliG region. We speculate that this flexibility plays a crucial role in facilitating the interaction between the cytoplasmic region of PomA and FliG to generate force.

**IMPORTANCE:** MS-ring is the initial structure to be assembled in flagellar motors. It comprises a complex two-ring (M and S) structure composed of 34 FliF molecules. We prepared a FliF–FliG fusion protein, which is directly involved in force generation. We observed it enabled the efficient formation of the MS-ring. The FliG portion that usually comprises the C-ring along with FliM and FliN displayed high flexibility likely due to the lack of FliM and FliN in the fusion protein. This study represents a significant milestone in the *in vitro* reconstruction of Na^+^-driven motor complexes.

## INTRODUCTION

The marine bacterium *Vibrio alginolyticus* possesses a single flagellum for propulsion in an aqueous environment, which functions as a motility organ at the cell pole (1). The flagellum is rotated by an ion-driven force of Na^+^ flow. A motor or force-generating complex at the base of the flagella acts as a power engine. The motor comprises a stator, ion energy conversion unit, and rotor that generates torque through interaction with the stator. Earlier, the flagellar motor thought to be the basal body of the flagellar structure; however, a new structure has been discovered at the base of the flagella in the membrane (2). Additionally, adenosine triphosphate (ATP) had been considered the energy source for the flagellar motor. In the basal body of gram-negative bacteria, the LP-ring corresponds to the outer membrane/peptidoglycan layer, whereas the MS-ring corresponds to the inner membrane (2). An axial structure at the center known as the rod connects the ring structures (Fig. 1). The MS-ring, which possesses a double-ring structure, is the main motor component that generates torque, and FliF is at least a constituent protein of the M-ring (3). The earlier hypothesis was that the M- and S-rings were composed of distinct proteins and rotation occurred between these two ring structures (4). Subsequently, the MS-ring was revealed to be a complex two-ring structure composed solely of the FliF protein (5). FliG, FliM, and FliN form a C-ring just below the MS-ring (6–8). FliG is one of the most important proteins in the C-ring that directly generates torque, and the C-terminal charged residues of FliG interact with the charged residues of the stator composed of the motor membrane proteins MotA and MotB or PomA and PomB (9–12). Recently, a direct interaction between the stator and rotor has been detected (13).

**Fig. 1.**
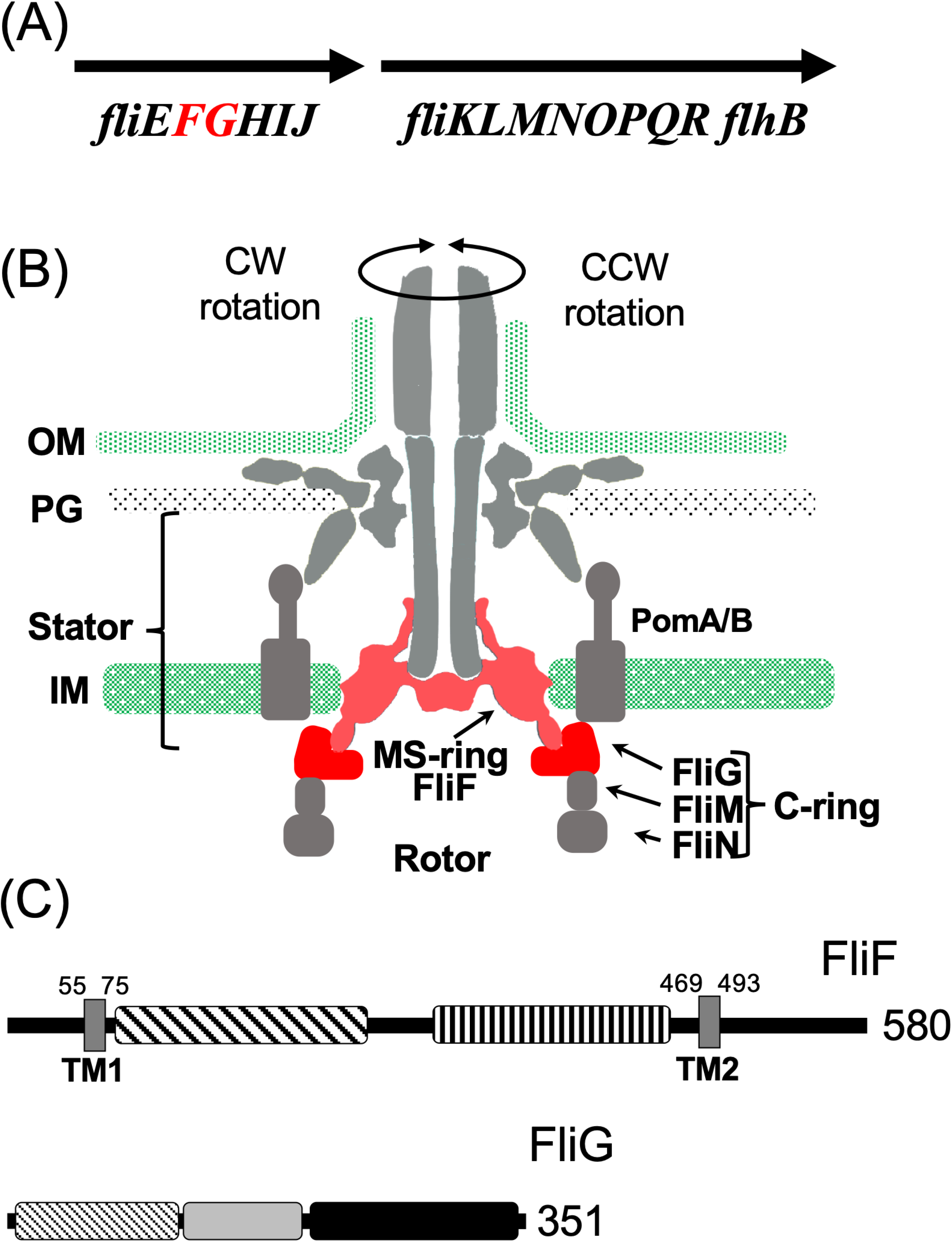
Flagellar motor structure and the component genes. (A) The genomic context of the flagellar genes around *fliF* and *fliG*. (B) The longitudinal section image of the flagellar motor. The C-ring is composed of FliG, FliM, and FliN. The MS-ring is composed of FliF. The stator is composed of PomA and PomB. FliF and FliG are indicated in red. (C) The primary structure of FliF and FliG. The distinctive regions of the protein are represented by the squares painted with different patterns. TM: transmembrane region. The numbers indicate the amino acid residues.

The construction scheme for flagellar motors was proposed more than 40 years ago after isolating intermediate structures from flagellar-deficient mutants (14); this basic structure formation model is still used today (1). However, the mechanism underlying the initiation of flagellar structure assembly and control of the assembly pathway remains unclear. The MS-ring, which is embedded in the membrane at the flagellar base, is probably the first component produced during flagellar formation. The overexpression of *Salmonella* FliF in *E. coli* results in the formation of numerous MS-rings (5). However, when *Vibrio* FliF was overexpressed in *E. coli*, the MS-ring was not isolated and most proteins were purified as soluble protein complexes (15). We showed that this complex had the ability to interact with FliG, which is a component protein of the C-ring, and the N-terminal region of FliG interacted with the C-terminal region of FliF.

As mentioned earlier, the *Salmonella* FliF forms an M-ring structure when overexpressed in *E. coli*. Therefore, this system was utilized to purify the MS-ring, and its structure was elucidated using cryo-electron microscopy (16). However, the M-ring could not be resolved at the atomic level, probably because of its flexible structure. FliF weighs approximately 64 kDa. It has two transmembrane proteins made of 34 molecules that are assembled into an MS-ring within the membrane. Based on the FliF crystal structure, the M-ring region was inferred to be in proximity to the C-ring except for the corresponding portion in the transmembrane region (17). As mentioned earlier, *Vibrio* FliF does not form the MS-ring structure even when overexpressed in *E. coli*, but co-expression with FliG or FlhF promotes MS-ring formation using FliF in *E. coli* (18). FlhF is an essential factor in the generation of polar flagella and is involved in determining the number of flagella (19, 20). No other genes homologous to *flhF* have been reported in *E. coli* or *Salmonella*. Deletion of *flhF* results in the inability of *Vibrio* to form polar flagella (19), and deletion of the *sflA* gene (21), which encodes a transmembrane protein containing the cytoplasmic DnaJ motif and the periplasmic TRP motif, results in FlhFG-deficient cells forming flagella from any location outside the polar region (22, 23). Based on these experimental facts, SflA has been speculated to inhibit FliF from forming an MS-ring (Fig. S1).

In this study, based on the report of the functional FliF–FliG fusion protein in *S. enterica* (7), we successfully cloned the *V. alginolyticus fliF–fliG* fusion gene into the pCold vector and overexpressed it in *E. coli*. We aimed to purify the fusion protein and investigate its function and structure.

## RESULTS

### Construction and expression of *fliFG* fusion gene

The analysis of the transcriptional regulation of the polar flagellar gene in *Vibrio* showed that the gene encoding the MS-ring constituent of the FliF protein belongs to an operon containing the genes for flagellar basal body proteins (24–27) (Fig. 1A). The *fliE* gene, which encodes the rod protein located in the MS-ring structure in the basal body of *Salmonella* (28, 29), is the first gene of this polycistronic operon. The *fliF* and *fliG* genes are the second and third genes in the operon, respectively. This indicates that FliE is produced prior to FliF and FliG proteins.

The molecular weights of FliF and FliG were approximately 64 kDa and 39 kDa, respectively (Fig. 1C). The C-terminal coding region of FliF and N-terminal coding region of FliG overlapped. Therefore, we induced a deletion of a single nucleotide preceding the start codon (ATG) of *fliG*. This resulted in the loss of two C-terminal residues (Asn579 and Gly580) of FliF and fusion with the initial Met of FliG. The resultant FliF–FliG fusion construct (FliFG) was incorporated with a His-tag (approximately 100 kDa) at the N-terminus to facilitate purification. Thus, FliFG protein expression was induced and the fusion protein was purified (Fig. 2). SDS-PAGE was performed, and the fusion protein was isolated between the 95 and 130 kDa markers. Furthermore, FliFG protein showed reactivity against both FliF and FliG antibodies. Cells housing the fusion construct were cultured, harvested, subjected to ultrasonic lysis, centrifuged at low speed, and subsequently fractionated into membrane and cytoplasmic fractions through ultracentrifugation. The FliFG fusion protein was separated into cytosolic and membrane fractions, similar to when FliF alone was expressed. The membrane fraction was treated with detergents, and the FliFG fusion protein was recovered as the ultracentrifuged precipitate from the solubilized membrane fraction. Analysis of this fraction using electron microscopy revealed a ring structure as shown in the following section. The cytoplasmic and membrane fractions were purified using Hi-Trap TALON with a His tag (Fig. S2). The peak fraction was then concentrated and subjected to gel filtration. FliFG (approximately 350 kDa), which was derived from the cytoplasmic fraction, was separated using gel filtration (Fig. 3A). In contrast, the membrane fractions solubilized using surfactants were separated at a very large molecular weight near the void (Fig. 3B). This void peak fraction was further separated using SDS-PAGE and the FliFG fusion protein was detected using coomassie brilliant blue (CBB).

**Fig. 2.**
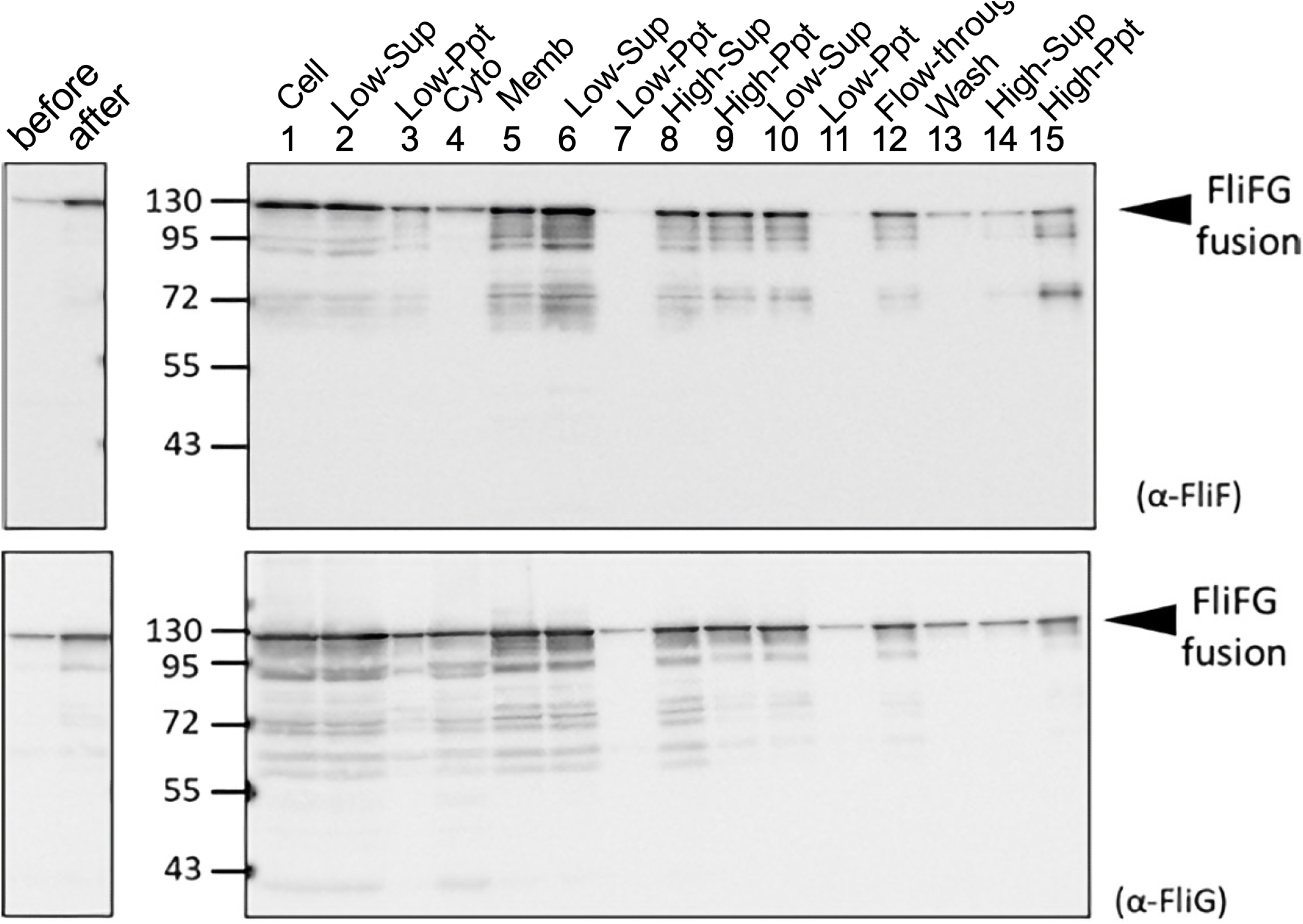
Purification of FliFG fusion protein. *E. coli* cells expressing FliFG fusion protein were harvested and suspended in the buffer. The cells (1: Cell) were sonicated, and the pellet (3: Low-Ppt) and supernatant (2: Low-Sup) were obtained using low-speed centrifugation. The supernatant was centrifuged at high speed to obtain the supernatant (4: Cyto) and the pellet (5: Memb). The membrane fraction was solubilized using a detergent, and the supernatant (6) and pellet (7) were obtained using low-speed centrifugation. The obtained supernatant (6) was centrifuged at high speed to obtain the supernatant (8) and the pellet (9). This pellet of (9) was suspended and centrifuged again at low speed to further separate the supernatant (10) and the pellet (11). The supernatant (10) was loaded in the column for his-tag. The flow-through (12) was recovered, washed with buffer (13), and eluted with imidazole. The eluted fraction was centrifuged at high speed to separate the supernatant (14) and the pellet (15). The proteins were separated using SDS-PAGE and detected using immunoblotting using anti-FliF (A) and anti-FliG (B) antibodies.

**Fig. 3.**
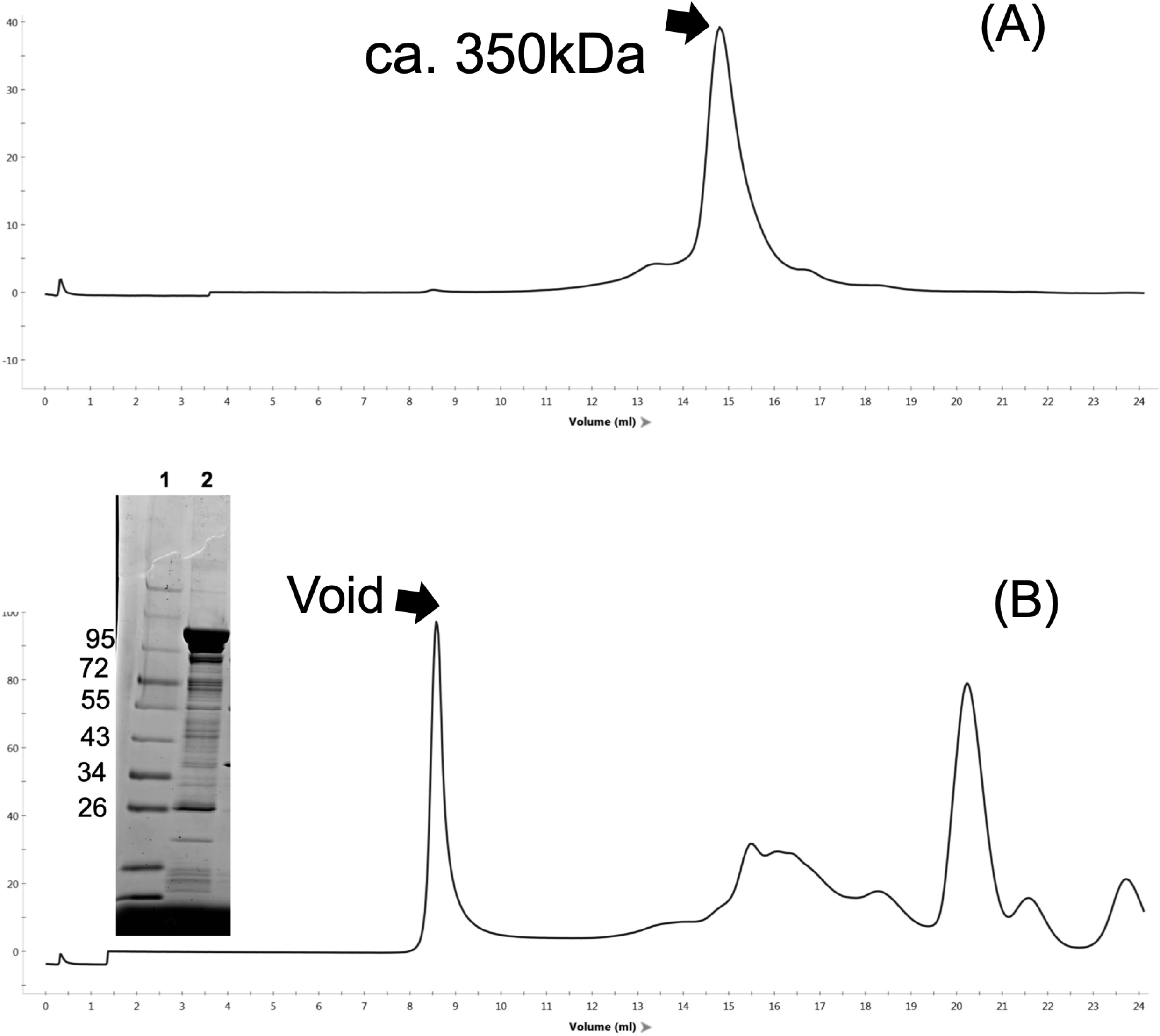
Elution profile of gel filtration. The protein samples (0.1 mL) of the cytoplasmic fraction of C (A) and the membrane fraction of CP (B) were run in a Superose 6 10/300 column with 20TN150 containing 1 mM MgCl_2_ and 0.05% dodecyl maltoside (DDM) at 0.5 mL/min flow rate. The proteins, 1: molecular weight markers, and 2: void fraction, were separated using SDS-PAGE and stained with CBB are shown in the inset of (B).

### Ring structure formation by the FliFG fusion protein

The precipitate obtained through high-speed centrifugation was viewed using electron microscopy with negative staining (Fig. 4, Fig. S3). This observation qualitatively showed that more ring formation in the purified precipitate from bacteria expressing the FliFG fusion protein than from those expressing FliF alone or co-expressing FliF and FliG (18). The FliFG fusion protein presented a ring structure (approximately 30 nm in diameter) that was similar to the structure previously isolated by expressing FliF without fusion with FliG. Some structures that were approximately 50 nm in diameter were discerned, albeit less distinctly, in the ring composed of the FliFG fusion protein around the well-defined ring structure.

**Fig. 4.**
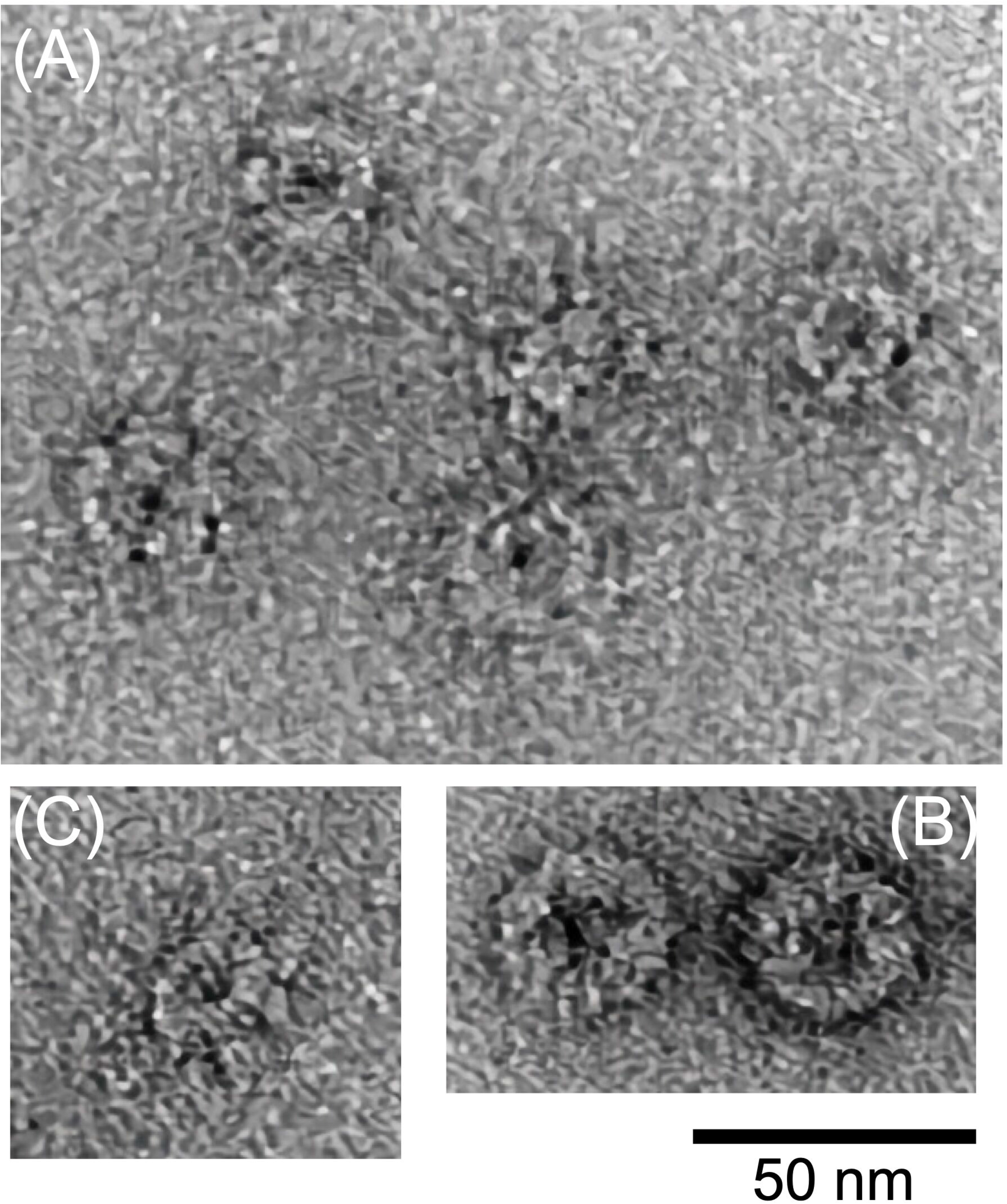
Electron microscopic observation of MS-ring composed of FliFG fusion proteins. The scale bars; 50 nm.

### HS-AFM observation of FliFG fusion protein

We used high-speed atomic force microscopy (HS-AFM) to observe the FliFG fusion protein that was separated using gel filtration near the void and the approximately 350 kDa protein that was absorbed onto a mica substrate using a solution containing detergent. The latter did not show any ring structures, and only a filamentous structure was observed (Fig. S4). The void sample showed a ring structure approximately 30 nm in diameter and approximately 20 nm in height, as shown in Fig. 5A. A magnified image (Fig. 5B) shows a vague structure within the area surrounded by a dashed line around a distinct ring structure that was approximately 30 nm in diameter. Since the vague structure is approximately 50 nm in diameter, this would correspond to the ring that was approximately 50 nm in diameter in the electron microscopy image. Vague structures were not observed in the FliF ring without fusion with FliG, as shown in Fig. 5C, indicating that the vague structure was derived from FliG. Fig. 5D shows successive images clipped from a high-speed AFM movie taken at an imaging rate of 0.15 s/frame with contrast enhancement of the vague structure. The HS-AFM movie (Movie S1) reveals that the vague structure is highly flexible, indicating that FliG in the FliFG protein is unstructured or at least present without FliM or FliN, which generally combine with FliG to form the C-ring.

**Fig. 5.**
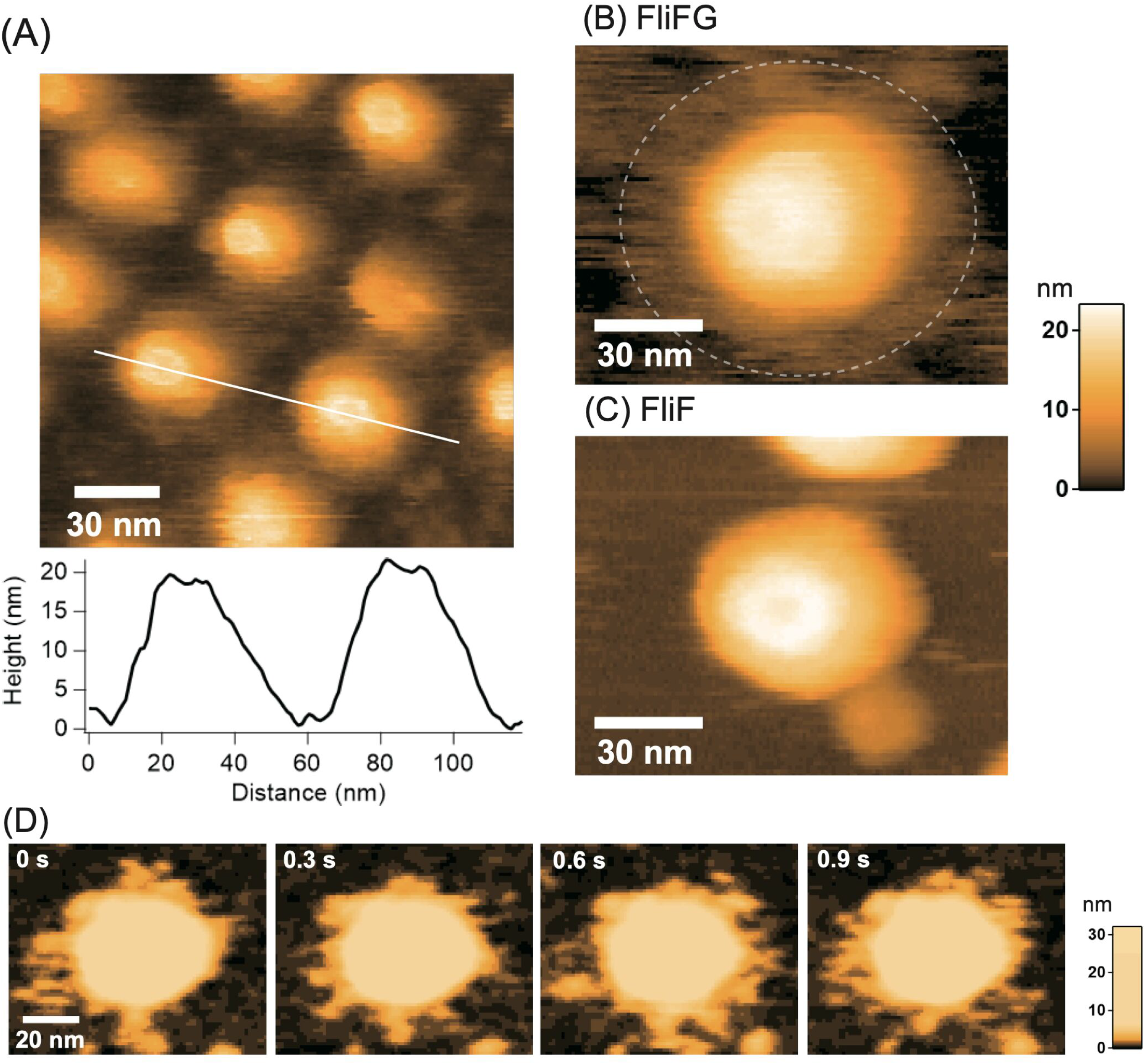
High-speed atomic force microscopy (HS-AFM) images of the purified *Vibrio* MS-ring. (A) HS-AFM image of MS-ring comprising FliFG fusion proteins and the cross-sectional profile along the white line on the image. (B) Magnified image of MS-ring composed of the FliFG fusion proteins. (C) HS-AFM image of MS-ring composed of the FliF proteins. (D) Clipped images of the MS-ring composed of FliFG fusion proteins with enhanced contrast of the lower part. Imaging rate: 0.15 s/frame.

## DISCUSSION

The flagellar structure is one of the most prominent architectures in bacteria. The smallest biological rotary motor for locomotion is present at the base of the flagellum in the bacterial membrane (1). The construction of this large and complicated structure commences following the formation of the foundational structure—the MS-ring. Thus, the MS-ring is speculated to be the initial flagellar structure to be formed. However, to complete the flagellar structure, the C-ring must be constructed directly beneath the MS-ring structure and a flagellar-specific transport apparatus needs to be constructed between the MS- and C-rings (8). Two different models for the initiation of flagellar formation have been presented in different studies: 1) the transport apparatus is assembled following that of the MS-ring, and part of the C-ring is formed by FliF and FliG (30), and 2) the core of an export gate, which is a part of the transport apparatus consisting of FliPQR, is assembled on the membrane, and the MS-ring structure is assembled around the FliPQR complex as the core (31, 32). However, when FliF was overexpressed, the MS-ring was formed using FliF alone (6, 18), suggesting that the FliPQR complex and FliG may function as catalysts for ring formation.

The hook-basal body was purified from *V. alginolyticus* using Triton X-100 detergent, resulting in the detection of 11 bands with approximate molecular weights of 62, 50, 47, 46, 40, 33, 32, 31, 28, 26, and 25 kDa (33). FlgI, MotX, and MotY were assigned to the bands. However, FliG and FliM were not detected in the hook-basal body (34). Therefore, we changed the purification procedure and used a different detergent, CHAPS (3-[(3-chloramidopro-pyl)-dimethylammonio]-1-propanesulfonate), and FliG was detected in the hook-basal body. However, FliM was not detected using immunoblotting, and the C ring remained unobserved in the electron microscopy image. The FliM protein seems to dissociate easily from the basal body and MS-ring. The *Vibrio* FliF protein, which is a membrane protein with two transmembrane regions and produced at high levels in *E. coli*, was purified and detected in the soluble fraction. It was estimated to be an oligomer of approximately 700 kDa using an analytical size exclusion chromatography (15). This indicates that the protein was translated or produced without being recognized as a membrane protein. For a protein to be inserted into the membrane, the transmembrane hydrophobic region generally needs to be recognized by the SRP (signal recognition particle) protein in membrane protein transports (35). We hypothesized that (Fig. S1) FlhF recognizes FliF in the translocon machinery of the Sec system (18). We recently used a pull-down assay with the FliF fragment fused to GST to demonstrate that the N-terminal hydrophobic region interacts with FlhF (36).

We have shown that *Vibrio* MS-ring formation in *E. coli* was promoted by the co-production of FliG; however, the isolated MS-ring did not contain FliG (18). In contrast, MS-ring formation in *E. coli* was not affected by the deletion of the C-terminus of the FliF protein in the presence of FlhF; however, the defective FliF–FliG binding induced by this deletion abolishes flagellation and confers a nonmotile phenotype (37). In this study, a fusion protein connected with *Vibrio* FliF and FliG was found to tandemly and efficiently form MS-rings in *E. coli*. Similarly, FliF and FliG fusion proteins of *Salmonella* conferred flagellated and motile phenotypes, although the rotation direction was biased towards clockwise (CW) rotation (7, 38). Using these mutants, we showed that the flagellar switch protein FliG localizes to the cytoplasmic M-ring face of the basal body, and this fusion restricts the switching of flagellar rotation. During basal body isolation, the C-ring is easily lost, suggesting that FliM and FliN easily dissociate from the M-ring (39). The formation of the ring structure by the FliFG fusion proteins using residue coevolution to generate homodimer building blocks for ring assembly was predicted based on the X-ray crystal structures of proteins from other species and injectosome analogs (40). An FliF–FliG ring model of *S. typhimurium* and *H. pylori*, which was generated by translation and docking to the MS–C ring map from *S. typhimurium* (41), was proposed based on the structural analysis of FliF and FliG complexes (42, 43).

Recently, we observed the intact structure of the basal body with a C ring in *V. alginolyticus* using cryo-EM tomography and reported that the ring diameter of the FliG region is different in C rings constructed from CCW and CW mutant FliG (44). The isolated motor–hook complex or the basal body without the C-ring from *Salmonella* was subjected to cryo-EM single particle analysis for structural determination (45). The density map of the MS-ring showed that it has complicated symmetries and is composed of 34 β-collar, 34 RBM3, and 23 RBM2 FliF domains; however, the protein densities for the RBM1 domain and inner membrane-binding region in FliF could not be determined. Even in the structural analysis of the MS-ring composed of *Salmonella* FliF molecules expressed from a plasmid in *E. coli*, the density of the regions could not be determined using the cryo-EM single-particle analysis (16, 46). The cytoplasmic face of the M-ring is also difficult to determine at the atomic level. The region corresponding to the M-ring was determined using the crystal structure of RBM1 and RBM2 of *Aquifex aeolicus* FliF fragment (residue 58-213), which conformed with the cryo-EM structure of the MS-ring in *Salmonella* (17). However, the cytoplasmic face of the M-ring regions in the structure have not been completely elucidated probably because these regions are very flexible, as evidenced directly using HS-AFM in this study. The regions consisting of FliG and facing the M-ring also appeared to be very flexible. In addition to the intrinsic flexible structure, the FliG ring region or part of the C-ring structure seemed unable to form a stable ring structure without FliM or FliN, and electron microscopy did not show a clear ring structure in the purified base body of *Salmonella* (7, 41).

In a subsequent study, we plan to add FliM and FliN to *E. coli* cells to produce the FliFG fusion protein, construct the basal structure of MS-ring with C-ring, and reconstruct the motor with the stator complex in this system. The present study would provide the groundwork needed for the reconstruction of the whole *Vibrio* flagellar motor *in vitro* in *E. coli*.

## MATERIALS AND METHODS

### Bacterial strains and plasmids

The bacterial strains and plasmids used in this study are listed in Table S1. *E. coli* was cultured in LB broth [1% (w/v) bactotryptone, 0.5% (w/v) yeast extract, and 0.5% (w/v) NaCl]. Ampicillin was added at a final concentration of 100 μg/mL to the *E. coli* culture.

### Construction of the FliFG fusion mutant

As previously described, a one-step PCR-based method was used to clone the *fliF* and *fliG* genes into a pCold vector (47). To fuse *fliF* and *fliG*, a deletion was introduced at the desired position using the QuikChange site-directed mutagenesis method described by Stratagene (15).

### MS-ring purification

*E. coli* BL21(DE3) cells housing the pRO101 or its derivatives (for expression of FliF or its deletions) were inoculated from a frozen stock into a plate containing the appropriate antibiotics, and the colonies were inoculated into 3 mL of LB broth and cultured overnight with shaking at 37 °C. Then, 1 mL of the overnight culture was added to 50 mL of LB broth containing ampicillin and cultured at 37 °C with shaking until the OD_600_ value was approximately 0.5. Isopropyl β-D-1-thiogalactopyranoside (IPTG) (0.5 mM) was then added. The cells were subjected to cold shock by placing in ice-cold water for 20 min and cultured with shaking at 16 °C overnight. The cells were collected (4,600 × *g*, 10 min) and suspended in 30 mL of 20TK200 buffer (20 mM Tris–HCl [pH 8.0], 200 mM KCl) containing 1 mM ethylenediaminetetraacetic acid (EDTA). The cells were then sonicated (large probe, power=6, 60 s, three times, duty cycle 50%). The unbroken cells were removed through centrifugation at 10,000 × *g* for 10 min at 4 °C, and the cells were resuspended in 20 mL of 20TK200 containing 1 mM EDTA; the sonication process was repeated. The resulting supernatants were collected, and 2 mM and 20 mM of MgCl_2_ and imidazole, respectively, were added. Centrifugation was performed at 12,000 × *g* for 10 min at 4 °C, and the supernatant was further ultracentrifuged at 154,000 × *g* for 60 min at 4 °C. The supernatant and pellet were defined as the soluble cytoplasmic and membrane fractions, respectively. The precipitate of the membrane fraction was suspended in 20 mL of 20TK200 and shaken overnight in a cold room. Then, 2 mL of 10% dodecyl maltoside (DDM) solution or 1 mL of 10% Lauryl Maltose Neopentyl Glycol (LMNG) and 100 μL of 2 M imidazole were added and centrifuged at 10,000 × *g* for 10 min. The supernatant was used for Ni-affinity purification using a His-tag.

The sample was loaded onto a HiTrap TALON column (5 mL, GE Healthcare) connected to an ÄKTAprime system (GE Healthcare) and washed with 20TK200 buffer containing 0.05% DDM or 0.01% LMNG and 10 mM imidazole. The proteins were eluted and collected using an imidazole linear gradient of 10–300 mM. The peak fractions were concentrated through centrifuging in a 100 K Amicon device (Millipore). The samples were subjected to size-exclusion chromatography using Superose 6 column 10/300 with 20TN150 containing 1 mM MgCl_2_ and 0.05% DDM or 20TK100 with 0.0025% LMNG at a flow rate of 0.5 mL/min.

### Observation using electron microscopy

The purified MS-ring was observed after negative staining under an electron microscope. After hydrophilizing the carbon-coated copper grids, 2.5 μL of the sample solution was placed on the grids and stained with 2% (w/v) phosphotungstic acid (pH 7.2). The grid was observed under a transmission electron microscope (JEM-1010, JEOL) at 100 kV.

### High-speed-atomic-force-microscope (HS-AFM) observation and image analysis

HS-AFM imaging was performed using a laboratory-built HS-AFM operated in tapping mode (48). The AFM cantilever was AC7 purchased from Olympus (spring constant approximately 0.2 N/m and resonance frequency approximately 700 kHz in a solution). The AFM tip was fabricated by forming carbon pillars on the cantilever end using electron beam deposition, followed by plasma etching in an Ar environment to obtain the small end necessary for high-resolution imaging. The free oscillation amplitude of the cantilever for feedback control in HS-AFM imaging was reduced from approximately 2 nm to 1.5 nm.

The FliFG and FliG proteins were deposited on a bare mica substrate for HS-AFM imaging. After 5 min of incubation, the residual proteins were washed off using the observation buffer (20TK100 containing 0.0025% LMNG). HS-AFM imaging was performed in the solution at room temperature.

## Acknowledgments

We thank Kimika Maki for technical support with electron microscopy and Ryo Ogawa for plasmid construction of pRO301. This work was partially supported by JSPS KAKENHI Grant Numbers 21H00393, 21H01772, 22K18943 (to T.U.), and 20H03220 (to M.H.).

